# Cholesterol and ceramide facilitate SARS-CoV-2 Spike protein-mediated membrane fusion

**DOI:** 10.1101/2022.12.16.520599

**Authors:** Kristina Niort, Julia Dancourt, Erwan Boedec, Zahra Al Amir Dache, Grégory Lavieu, David Tareste

## Abstract

SARS-CoV-2 entry into host cells is mediated by the Spike (S) protein of the viral envelope. The S protein is composed of two subunits: S1 that induces binding to the host cell *via* its interaction with the ACE2 receptor of the cell surface and S2 that triggers fusion between viral and cellular membranes. Fusion by S2 depends on its heptad repeat domains that bring membranes close together, and its fusion peptide (FP) that interacts with and perturb the membrane structure to trigger fusion. Recent studies suggest that cholesterol and ceramide lipids from the cell surface may facilitate SARS-CoV-2 entry into host cells, but their exact mode of action remains unknown. We have used a combination of *in vitro* liposome-liposome and *in situ* cell-cell fusion assays to study the lipid determinants of S-mediated membrane fusion. We found that cholesterol and ceramide both facilitated fusion, suggesting that targeting lipids could be effective against SARS-CoV-2. As proof of concept, we examined the effect of chlorpromazine (CPZ), an antipsychotic drug known to perturb membrane structure. We found that CPZ inhibited S-mediated membrane fusion and thus potentially SARS-CoV-2 entry.

## Introduction

Cellular infection by SARS-CoV-2 starts with the fusion of its lipid envelope with the host cell membrane that leads to the delivery of its genomic RNA into the cell cytoplasm, production of new viral proteins by the host cell, and assembly/release of newly formed viruses. Such fusion event can occur directly with the cell plasma membrane or with the endosomal membrane following endocytosis of SARS-CoV-2. Because it occurs early in the viral replication cycle, this early fusion step is an attractive target for the development of drugs and vaccines able to block virus entry^1^.

SARS-CoV-2 binding and fusion with the cell membrane is mediated by the Spike (S) protein, a class I viral fusion protein composed of two subunits, S1 and S2. The S1 subunit is involved in binding to the host cell plasma membrane *via* its interaction with the human angiotensin-converting enzyme (ACE2) receptor of the cell surface^2–4^, whereas the S2 subunit mediates the fusion of SARS-CoV-2 lipid envelope with the host cell membrane^5^. For the S2 subunit to be available for fusion, the S protein must be cleaved at the S1/S2 interface by cellular proteases after S1 binding to ACE2. These proteases can be either from the cell surface, *e*.*g*. the membrane-anchored serine protease TMPRSS2^6^, when fusion occurs directly at the plasma membrane, or from endosomes, *e*.*g*. the cysteine protease cathepsin L^7^, when fusion occurs at the endosomal membrane after endocytosis of the viral particle. Several lines of evidence obtained with SARS-CoV-2 or the closely related coronaviruses SARS-CoV and MERS-CoV suggest that direct fusion with the plasma membrane is the preferred route, notably during infection of respiratory cells^1^. S protein produced by infected cells can also be directly transferred to the cell surface^8^, leading to potential fusion with neighboring uninfected cells, which is believed to play a key role in SARS-CoV-2 pathogenicity^9^. The S2 subunit of the SARS-CoV-2 S protein possesses an N-terminal fusion peptide (FP), followed by two heptad repeat domains (HR1 and HR2), and a transmembrane domain (TMD) that anchors the S protein into the viral membrane. Fusion by S2 starts with the insertion of its FP into the target cell membrane to establish a molecular bridge consisting of a three-helix coiled-coil complex composed of its HR1 and HR2 domains. S2 then folds back onto itself, in the form of a six-helix coiled-coil hairpin complex of the HR1 and HR2 domains, which brings the FP and TMD in molecular proximity along with the viral and cellular membranes in which they reside. This close membrane apposition combined with lipid bilayer destabilization produced by membrane insertion of the FP leads to fusion. Isolated FPs have been very useful in elucidating the mechanisms of viral fusion because (i) they can themselves induce fusion, and (ii) single mutations within FPs can lead to complete loss of viral fusion and infection^10,11^. FPs usually consist of about 20 amino acids, mostly hydrophobic, and are highly conserved within a given viral family. Contrary to other type I viral fusion proteins, such as those of HIV and influenza viruses which possess a single FP, several potential FPs with membrane interacting and/or membrane fusion activity were identified within the S2 subunit of coronaviruses S protein^12^. The region FP1 (residues 816-833 in SARS-CoV-2) is however believed to be the functional FP based on its high sequence conservation (particularly among MERS-CoV, SARS-CoV and SARS-CoV-2) and its location immediately downstream of the S2’ cleavage site that is known to be critical for coronaviruses fusion^6,10,13,14^ (Fig. 1A). In the case of SARS-CoV, FP1 was shown to induce membrane fusion and some mutations that abolished FP1-mediated liposome fusion *in vitro* also affected cell-cell fusion and viral infection *in situ*^10^. The region just C-terminal of FP1 (residues 834-855 in SARS-CoV-2, called FP2) was proposed to complement FP1 to form an extended fusion peptide (FP1-FP2) with higher membrane affinity^15^. Recent experimental and computational structural studies showed that the FP1 of SARS-CoV-2 inserts deeply into lipid bilayer structures, whereas its FP2 stays around the bilayer surface^16–18^, suggesting strong membrane perturbing effect and thus fusion activity by FP1. But direct evidence that the FP1 of SARS-CoV-2 can induce membrane fusion is still missing.

**Figure 1.**
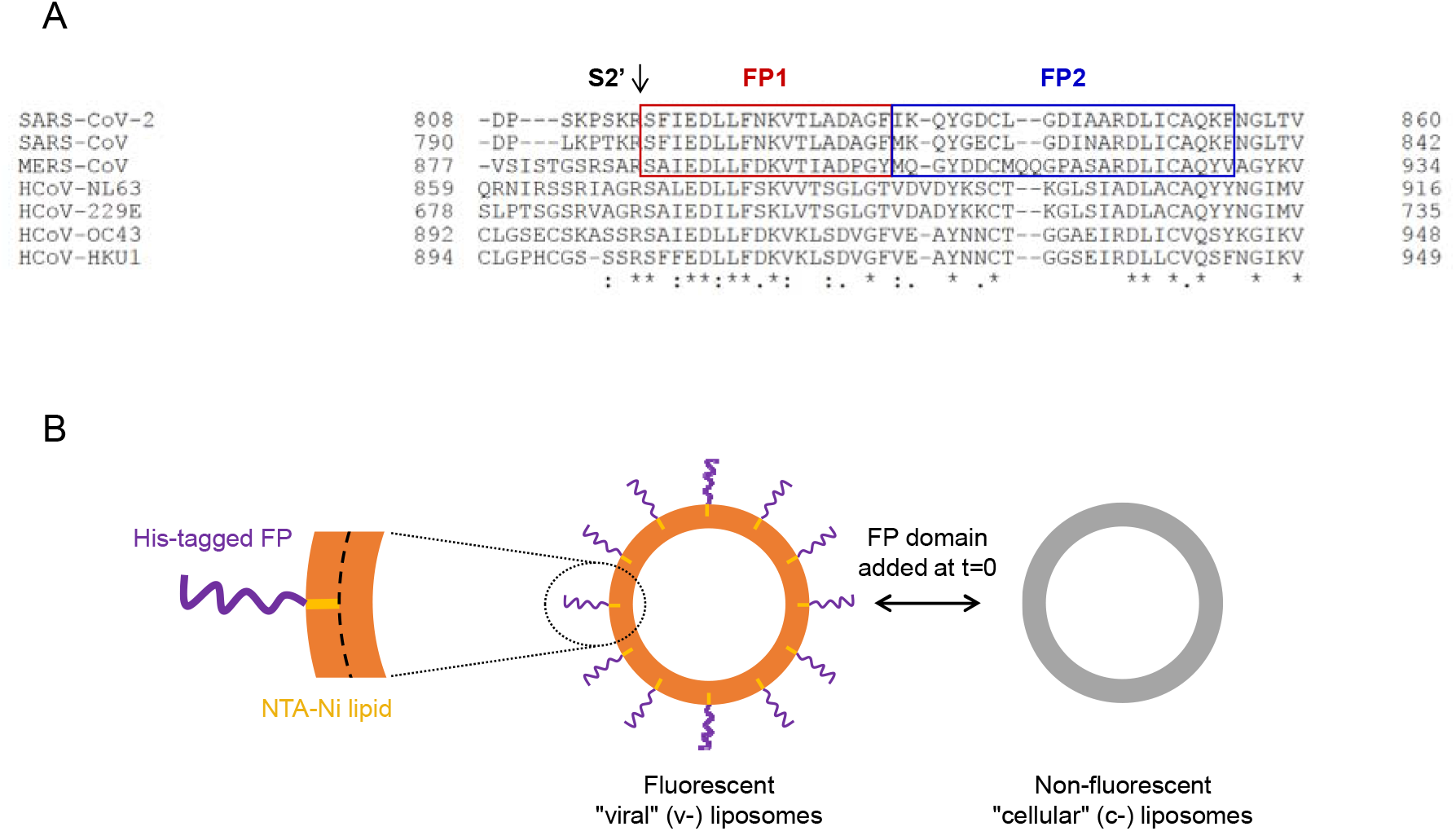
(A) Sequence alignment of Spike proteins from all human coronaviruses in the region immediately following the S2’ cleavage site known to be critical for SARS-CoV-2 fusion. The fusion peptides FP1 (red) and FP2 (blue) are highly conserved notably among MERS-CoV, SARS-CoV and SARS-CoV-2. (B) Experimental set-up used to study the capacity of SARS-CoV-2 fusion peptides to mediate membrane fusion *in vitro*. Fusion peptides with a C-terminal His_6_ tag were reconstituted at t=0 of the assay into fluorescently labeled liposomes mimicking the viral envelope (v-liposomes) by binding to lipids having an NTA-Ni headgroup. Fusion was monitored between v-liposomes and unlabeled peptide-free liposomes mimicking the cellular membrane (c-liposomes) using a FRET-based lipid mixing assay^29^.

The activity of viral FPs also depends on the lipid composition of the fusing membranes^19^. The FP of SARS-CoV was found to bind stronger and to penetrate deeper into membranes containing cholesterol (CHOL)^20^. Interestingly, a recent study showed that the enzyme cholesterol 25[hydroxylase could inhibit infection by MERS-CoV, SARS-CoV and SARS-CoV-2 by inducing depletion of plasma membrane CHOL, thus blocking viral fusion at the cell surface^21^. Along the same lines, generation of ceramide (CER) membrane domains at the cell surface upon cleavage of sphingomyelin by the acid sphingomyelinase was found to facilitate SARS-CoV-2 entry into cells^22^. This suggests that repurposing drugs which can modify the structure or lipid composition of the host cell membrane could be an effective treatment against SARS-CoV-2 infection. In this regard, antipsychotic (AP) drugs are interesting candidates that were proposed to exert some antiviral activity against coronaviruses^23–25^. Because of their amphiphilic property and planar geometry, AP drugs are known to intercalate into lipid bilayers and thus modify their biophysical properties^26^, which can in turn impact the ability of these lipid bilayers to fuse.

In this paper, we have investigated *in vitro* with a fluorescence resonance energy transfer (FRET)-based liposome fusion assay the capacity of FP1 and FP2 from SARS-CoV-2 S protein to fuse membranes of various lipid compositions (including or not CHOL and CER). We have also tested the ability of the AP chlorpromazine (CPZ) to inhibit FP-mediated membrane fusion. The effect of lipid composition and CPZ addition was also examined *in situ* on full-length S protein-mediated membrane fusion measured by a nanoluciferase-based cell fusion assay.

## Results

The fusion activity of FP1 and FP2 from SARS-CoV-2 S protein was examined *in vitro* through their capacity to mediate the fusion between two liposome populations mimicking viral and cellular membranes, respectively. We used synthetic peptides with a C-terminal His_6_ tag allowing their chemical coupling to reactive NTA-Ni lipids included in the liposome membrane (Fig. 1B). Similar lipid-anchorage strategy has proven successful in recapitulating SNARE-mediated membrane fusion *in vitro*^27^, and more recently in revealing the role of the heptad repeat domains of Mitofusin in mitochondrial membrane docking and fusion^28^. Fusion was measured using a FRET-based lipid mixing assay^29^. One population of liposomes (the fluorescent donor liposomes mimicking viral membranes, v-liposomes) had NTA-Ni lipids in their bilayer, whereas the other population (the non-fluorescent acceptor liposomes mimicking cellular membranes, c-liposomes) did not have any NTA-Ni lipid (Fig. 1B). This allowed FP1 and FP2 to get exclusively anchored onto the v-liposome surface. FP1 and FP2 were added at t=0 of the lipid mixing assay and their capacity to mediate fusion between v- and c-liposomes was monitored for 90 min.

We started with v- and c-liposomes composed exclusively of phosphatidylcholine (PC) lipid, one of the main lipids of biological membranes. FP1 induced robust fusion between PC v- and c-liposomes, whereas no significant fusion was observed with FP2 under the same experimental conditions (Fig. 2). In addition, no fusion was measured when FP1 was added to v-liposomes lacking NTA-Ni lipids (Fig. S1), indicating that FP1 needed to be membrane-anchored to induce lipid mixing in our system.

**Figure 2.**
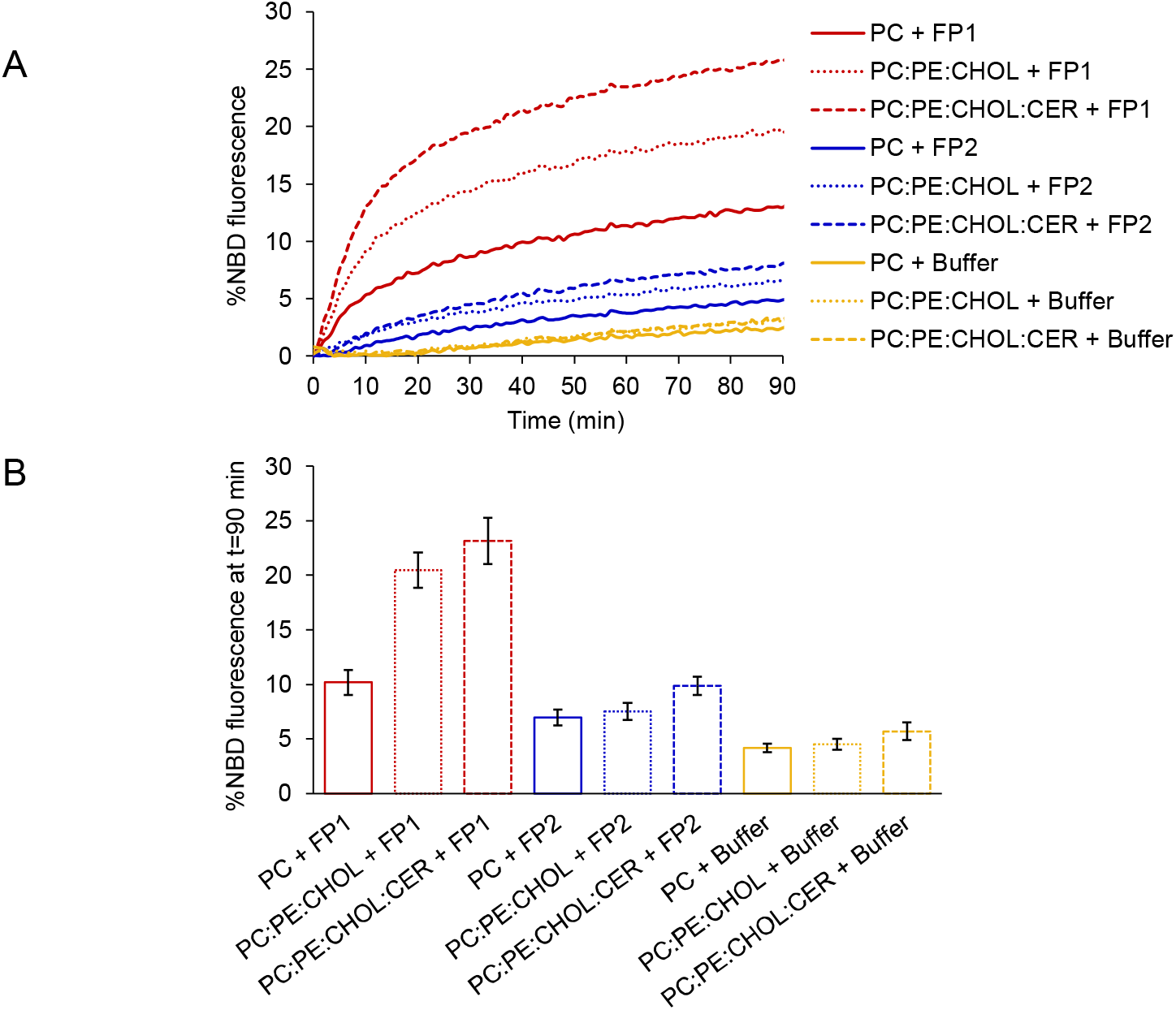
(A) Representative FRET-based lipid mixing experiments between v-liposomes (composed of 92 mol% PC, 5 mol% NTA-Ni, 1.5 mol% NBD and 1.5 mol% Rho) and c-liposomes of various lipid compositions (including or not 10 mol% PE and 30 mol% CHOL or 10 mol% PE, 30 mol% CHOL and 20 mol% CER at the expense of PC) in the absence (yellow) or presence of FP1 (red) or FP2 (blue) added at t=0 (500 μM of lipids and 25 μM of peptides). FP1 induced efficient lipid mixing between v- and c-liposomes exclusively composed of PC lipid. Fusion was strongly activated when the c-liposome membrane contained PE and CHOL or PE, CHOL and CER. No significant lipid mixing was measured under the same conditions with the FP2 peptide, or when the fusion peptides were not lipid-anchored (v-liposomes devoid of NTA-Ni lipids; see Fig. S1). (B) Average extent of lipid mixing after 90 min (n=4-9 independent experiments; error bars are standard errors).

To study the effect of cellular membrane lipid composition on FP1-mediated fusion, we measured the fusion between v-liposomes composed exclusively of PC lipids and c-liposomes including (in addition to PC) CHOL and CER lipids that were recently proposed to facilitate SARS-CoV-2 infection^21,22^. We also added phosphatidylethanolamine (PE) lipids in the c-liposome membrane to better mimic the lipid composition of the outer leaflet of mammalian plasma membranes^30^. Having PE and CHOL together in the c-liposome membrane dramatically increased FP1-mediated fusion (Fig. 2 and 3), whereas addition of either PE or CHOL alone had no significant effect on fusion (Fig. S2). Including CER in the c-liposome membrane (in addition to PC, PE and CHOL) further increased fusion mediated by FP1 (Fig. 2 and 3). For all lipid compositions tested, FP1 needed to be anchored to the v-liposome membrane to induce fusion with c-liposomes (Fig. S1).

**Figure 3.**
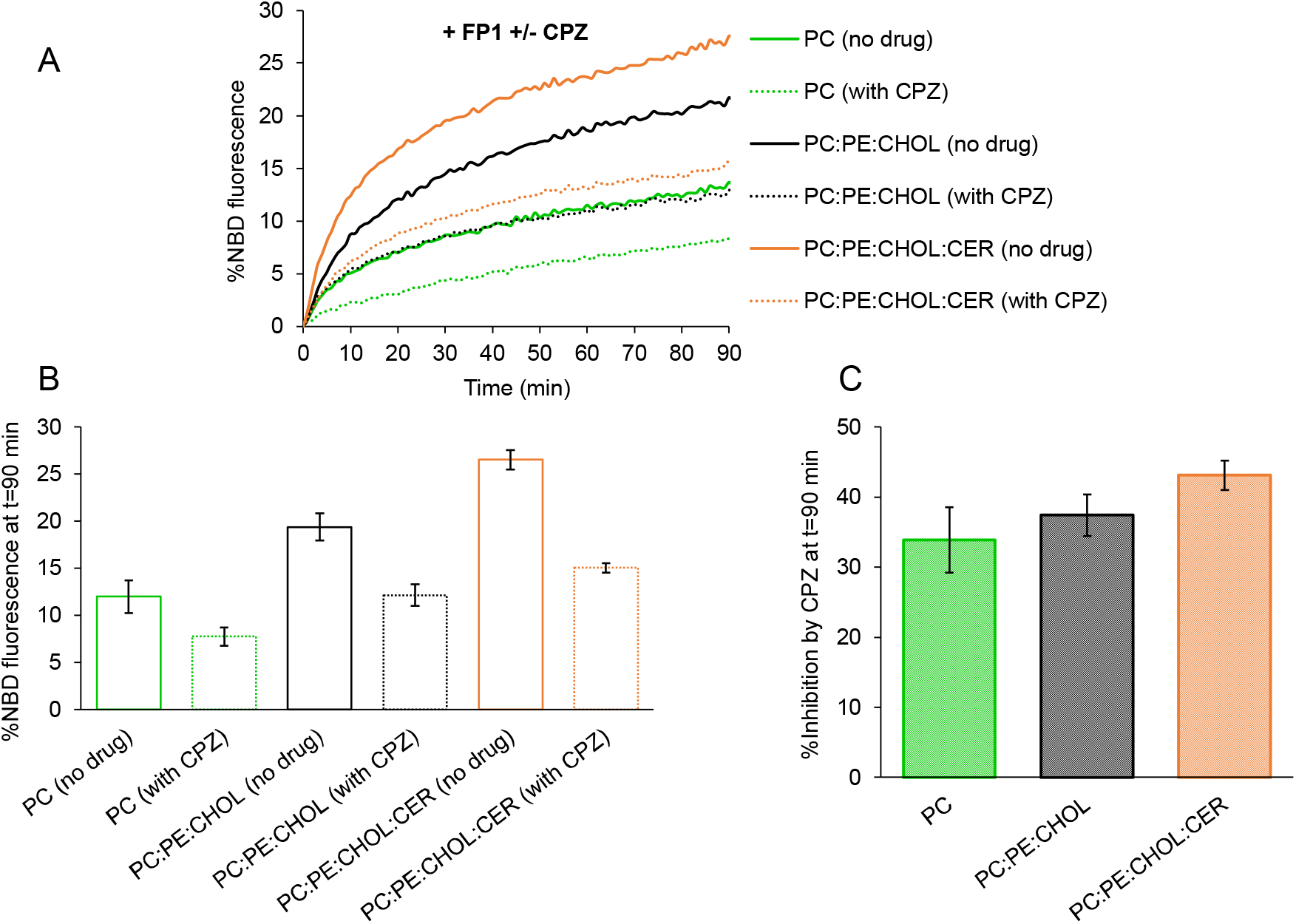
(A) Representative FRET-based lipid mixing experiments between v-liposomes (composed of 92 mol% PC, 5 mol% NTA-Ni, 1.5 mol% NBD and 1.5 mol% Rho) and c-liposomes of various lipid compositions (including or not 10 mol% PE and 30 mol% CHOL or 10 mol% PE, 30 mol% CHOL and 20 mol% CER at the expense of PC) in the presence of FP1 and in the presence/absence of chlorpromazine (CPZ), both added at t=0 (500 μM of lipids, 25 μM of peptides and 50 μM of CPZ). All fusion experiments (with or without CPZ) were performed in a final DMSO concentration of 1% (v/v) in buffer H. CPZ strongly inhibited FP1-mediated fusion between v- and c-liposomes regardless of the c-liposome membrane lipid composition. (B) Average extent of lipid mixing after 90 min (n=4-6 independent experiments; error bars are standard errors). (C) Percentage of fusion inhibition by CPZ after 90 min.

Fusion mediated by FP2 slightly increased upon modification of the c-liposome membrane lipid composition but remained low and close to the fusion background observed in the absence of any fusion peptide (Fig. 2 and S2). Of note, having PE in the c-liposome membrane somewhat activated all fusion events whether induced or not by fusion peptides. This agrees well with the known role of PE lipids as facilitators of membrane fusion^31^. Because high membrane curvature is known to activate fusion^32–34^, we analyzed the size of c-liposomes as a function of their lipid composition by multi-angle dynamic light scattering. We found that the size distribution of c-liposomes was not significantly affected by their lipid composition (Fig. S3). Importantly, c-liposomes exclusively composed of PC lipid were overall smaller than c-liposomes of more complex lipid compositions. Strong FP1-mediated membrane fusion measured in the presence of PE and CHOL or PE, CHOL and CER is therefore not due to high membrane curvature of c-liposomes.

Next, we tested the effect of the AP drug CPZ, which has affinity for lipidic phases and was shown to perturb membrane structural properties^26,35^, on FP1-mediated membrane fusion. CPZ was solubilized in dimethyl sulfoxide (DMSO) and added at the very beginning of the lipid mixing assay, together with the fusion peptide. The final DMSO concentration was 1% (v/v) to ensure that it would not affect the biophysical properties of liposome membranes^36^. CPZ inhibited FP1-mediated fusion between v- and c-liposomes and induced a similar decrease of about 40% of the extent of fusion for all lipid compositions tested (Fig. 3). To check if this observation was specific to CPZ, we also examined the effect of the antidepressant (AD) drug Fluvoxamine (FLUV), which also has amphiphilic and membrane-binding properties^37^. Under the same experimental conditions, FLUV only had a modest inhibitory effect on FP1-mediated fusion (Fig. S4).

The extended fusion peptide composed of FP1 and FP2 (FP1-FP2) was previously shown to exhibit greater membrane activity than FP1 or FP2 alone^15^. We thus wanted to check if this translated into a greater ability of FP1-FP2 to induce membrane fusion. We chose to work with c-liposomes composed of PC, PE and CHOL as we observed it to be the optimal composition for FP1-dependent fusion. FP1-FP2 triggered efficient fusion between v- and c-liposomes and the extent of lipid mixing at the end of the reaction (about 20%) was similar to that measured with FP1 (Fig. 3 and S5). The drugs CPZ and FLUV also had the same effect on FP1- or FP1-FP2-induced membrane fusion. CPZ decreased fusion between v- and c-liposomes by about 30% and FLUV did not have any significant effect on fusion (Fig. S5). In our system, FP1 and FP1-FP2 thus displayed the same fusion activity and responded similarly to the addition of the AP CPZ or the AD FLUV.

To confirm the biological relevance of our *in vitro* data, we used a cell-based assay in which we could test the impact of drugs that perturb lipid metabolism to mimic CHOL and CER depletion in the cell plasma membrane. We performed *in situ* measurement of cell-cell fusion between (i) donor HeLa cells that stably expressed the first half of a split nanoluciferase (HiBit-Hsp70) in their cytosol with or without Flag-tagged S proteins on their surface and (ii) acceptor HeLa cells that stably expressed the second half of the split nanoluciferase (LgBit) in their cytosol with or without ACE2 receptors on their surface. Split nanoluciferase complementation occurs only in case of cell-cell fusion (Fig. 4A). We incubated donor and acceptor cells in the presence or absence of S proteins and ACE2 receptors to establish that both proteins were required for efficient fusion. We observed that luminescence activity was well above the background signal only when S and ACE2 were present on donor and acceptor cells, respectively (Fig. 4B). This value was set to 100%. Note that in the absence of overexpressed ACE2 on acceptor cells, the luminescence activity was about 15%. This is consistent with the very low endogenous level of ACE2 on HeLa cells, which was shown to be below that required for SARS-CoV-2 infection but not so low as to prevent S-mediated cell-cell fusion ^2,38^. We then performed the same experiments in the presence of Fumonisin B1, a well-known inhibitor of CER synthase that dramatically reduces the level of CER^39^. This led to a 50% decrease of the fusion signal (Fig. 4B). When we added MβCD to artificially deplete the plasma membrane of CHOL, this also led to a strong (70%) inhibition of cell-cell fusion. Interestingly, CPZ treatment showed a similar effect on cell-cell fusion (70% inhibition) as removing CHOL, and we could not detect any additive effect when CPZ was added together with MβCD. Note that donor and acceptor cells were incubated together for at least 16 hours to allow them to adhere before the drugs were added, which may have caused occasional cell-cell fusion. Nevertheless, the cell-cell fusion assay confirms the *in vitro* results and shows that CHOL and CER depletion, as well as CPZ addition, inhibit S-mediated membrane fusion.

**Figure 4.**
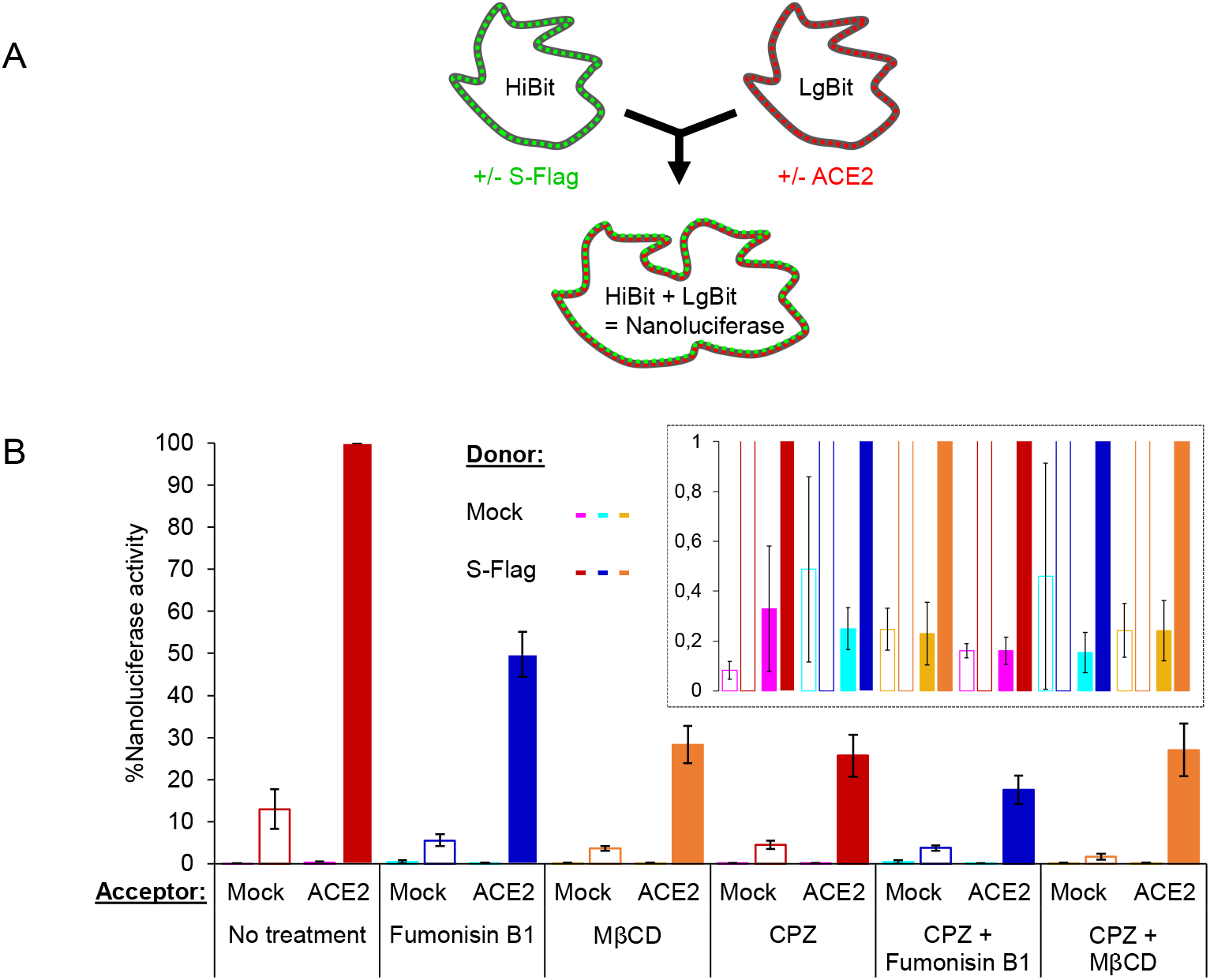
Effect of lipid composition and drugs on *in situ* cell-cell fusion. (A) Assay principle: HeLa cells stably expressing HiBit-Hsp70 and co-expressing or not C-terminally Flag-tagged wild-type Spike (S-Flag) were co-cultured with HeLa cells stably expressing LgBit and co-expressing or not ACE2. Cell-cell fusion triggers content mixing and nanoluciferase complementation. (B) Donor cells (HiBit-Hsp70 positive cells) co-expressing or not (mock) S-Flag were co-cultured with acceptor cells (LgBit positive cells) co-expressing or not (mock) ACE2. After 24 hours, Fumonisin B1 (20 μM) or MβCD (2 mM), and/or CPZ (10 μM) were added for 24 hours prior to reading nanoluciferase activity. The graph represents the percentage of nanoluciferase activity for each condition. Nanoluciferase activity resulting from content mixing between donor cells co-expressing S-Flag and acceptor cells co-expressing ACE2 was set to 100%. The inset shows the background nanoluciferase activity measured when donor cells do not co-express S-Flag (n=2 independent experiments in triplicates; error bars represent standard deviation of all replicates).

## Discussion

Our study shows that the fusion peptide FP1 of the S protein from SARS-CoV-2 mediates the fusion between liposomes to which it is membrane-anchored (v-liposomes mimicking the viral envelope) and protein-free liposomes (c-liposomes mimicking the cellular plasma membrane). On the contrary, the fusion peptide FP2 could not induce fusion between v- and c-liposomes in our system (Fig. 2 and S2). This is consistent with a recent structural study which found that FP1 becomes more structured in contact with lipid bilayers and inserts below the lipid headgroups, while FP2 remains unstructured and lies on the surface of lipid bilayers. It was then proposed that deeper membrane insertion in the case of FP1 should induce stronger perturbing effects on the lipid bilayer structure and thus favor fusion^16^. A structural study on FP1 and FP2 of the S protein from SARS-CoV found that FP1 and FP2 can cooperate and penetrate deeper into the lipid bilayer structure when they are together^15^. In our liposome fusion assay, FP1-FP2 displayed the same fusion activity as FP1 (Fig. 3 and S5). The synergy between FP1 and FP2 was thus not observed in our experiments with FP1 and FP2 from SARS-CoV-2 or at least it did not translate into stronger fusion activity.

We also observed that adding PE and CHOL into the c-liposome membrane to better recapitulate the lipid composition of the outer leaflet of mammalian plasma membranes strongly enhanced FP1-induced fusion between v- and c-liposomes (Fig. 2 and 3). In our cell-cell fusion assay, depletion of plasma membrane CHOL in the presence of MβCD induced a dramatic (∼70%) decrease of fusion between cells expressing S proteins and cells expressing ACE2 receptors (Fig. 4). *In vitro* liposome-liposome fusion and *in situ* cell-cell fusion assays therefore both show that CHOL facilitates S-mediated membrane fusion. This agrees well with two recent studies which found that efficient SARS-CoV-2 fusion with the cell membrane requires CHOL^9,21^.

Cellular infection by SARS-CoV-2 was also previously found to be facilitated by the presence of CER in the cell plasma membrane^22^. CER molecules are known to assemble into large hydrophobic gel-like domains in the outer leaflet of the cell plasma membrane that can serve to sequester specific surface proteins. It was proposed that such CER-enriched domains could trap ACE2 and TMPRSS2 to facilitate S protein binding and priming, respectively^40^. We hypothesized that CER-enriched domains, because of their high hydrophobicity, could also favor FP1 interaction with the cell membrane and thus membrane fusion. Addition of CER into the c-liposome membrane in fact increased FP1-mediated fusion between v- and c-liposomes *in vitro* (Fig. 2 and 3), and depletion of CER upon treatment of cells with Fumonisin B1 decreased S-mediated cell-cell fusion *in situ* (Fig. 4). CER could thus promote SARS-CoV-2 infection in two distinct ways: (i) by facilitating S protein binding/priming, and (ii) by directly activating the fusion step.

The observation that CHOL and CER facilitate S-mediated membrane fusion prompted us to investigate the effect of approved drugs that can modify membrane lipid composition and/or structure and could thus be repurposed to inhibit SARS-CoV-2 infection. We focused on the AP CPZ because of its previously proposed antiviral activity and its known effects on membrane biophysical properties. Several recent studies revealed that AP drugs could be effective in reducing SARS-CoV-2 infectivity. A screening assay based on morphological profiling of cells infected by SARS-CoV-2 identified two APs, domperidone and metoclopramide, exhibiting antiviral effects^23^. Another study searching for drugs targeting SARS-CoV-2 proteins interactome found that the AP haloperidol (HAL) displayed some antiviral activity^24^. Finally, CPZ was found to inhibit the replication of MERS-CoV, SARS-CoV and SARS-CoV-2 in cultured cells^25,41,42^. It was furthermore suggested that CPZ might block viral replication at an early entry stage, which could be the fusion step. Our *in vitro* and *in situ* fusion data both unambiguously show that addition of CPZ reduces S-mediated membrane fusion (Fig. 3 and 4). Interestingly, they also show that adding CPZ produces the same effect on fusion as depleting the membrane of CHOL (Fig. 2-4). CPZ was previously found to modify the lateral organization of CHOL-containing membranes and to perturb raft domains, which could be attributed to its capacity to compete with CHOL since CPZ and CHOL have a similar ring-like planar structure^26,35^. It is tempting to speculate that such competition between CPZ and CHOL could explain the inhibitory effect of CPZ on S-mediated membrane fusion observed in our study. This would also explain why CPZ had no effect on S-mediated cell-cell fusion when cells were treated with MβCD. *In vitro*, however, the inhibitory effect of CPZ persisted regardless of the lipid composition of the liposomes. This could be because (i) CPZ was also shown to increase lipid order in membranes or membrane regions lacking CHOL^26,35^ and (ii) *in vitro* systems, which are intrinsically simpler in terms of their lipid and protein composition, are more sensitive to perturbations of the membrane structure. Under the same experimental conditions, the AD drug FLUV, an amphiphilic ring-like molecule that also interacts with membranes but has little effect on their biophysical properties^37^, did not significantly alter FP1-mediated membrane fusion (Fig. S4). The inhibitory effect of CPZ on FP1-induced fusion could therefore result from its ability to increase lipid order and thus counterbalance the perturbing effect of FP1 on membrane structure.

So far, therapeutic strategies targeting the S2 fusion machinery mainly focused on the development of anti-fusogenic synthetic peptides mimicking the HR2 domain. Such peptides were shown to inhibit formation of the native HR1/HR2 six-helix coiled-coil hairpin complex, thus preventing viral fusion and infection by several human coronaviruses *in situ*, including MERS-CoV, SARS-CoV and SARS-CoV-2^43–45^. Furthermore, intranasal injection of a synthetic lipopeptide derived from HR2 was found to protect ferrets against infection by SARS-CoV-2^46^. These promising effects of anti-fusogenic synthetic HR2 peptides remain however to be evaluated in the context of human clinical trials. To circumvent the complex processes associated with the development of new drugs, an alternative strategy consists in repositioning existing approved drugs by testing their capacity to block SARS-CoV-2 fusion with target cells. Our study is in line with this approach and suggests that amphiphilic molecules with a planar shape such as CPZ could be effective inhibitors of SARS-CoV-2 infection due to their ability to modify the structure of membranes.

## Materials and Methods

### Chemicals

1-palmitoyl-2-oleoyl-*sn*-glycero-3-phosphocholine (PC), 1,2-dioleoyl-*sn*-glycero-3-phosphoethanolamine (PE), 1,2-dioleoyl-*sn*-glycero-3-phospho-L-serine-N-(7-nitro-2-1,3-benzoxadiazol-4-yl) (ammonium salt) (NBD), 1,2-dioleoyl-*sn*-glycero-3-phosphoethanolamine-N-(lissamine rhodamine B sulfonyl) (ammonium salt) (Rho) and 1,2-dioleoyl-sn-glycero-3-[(N-(5-amino-1-carboxypentyl)iminodiacetic acid)succinyl] (nickel salt) (NTA-Ni) were purchased from Avanti Polar Lipids as chloroform solutions. Cholesterol (CHOL) and N-stearoyl-D-erythro-sphingosine (CER) were purchased from Avanti Polar Lipids as powder and solubilized in analytical grade chloroform.

N-2-hydroxyethylpiperazine-N’-2-ethanesulfonic acid (HEPES, OmniPur grade), potassium hydroxide solution 47% (KOH 47%, EMSURE grade for analysis), potassium chloride (KCl, OmniPur grade), glycerol (Molecular Biology grade), n-octyl-β-D-glucopyranoside (β-OG, ≥ 98% GC), n-dodecyl β-D-maltoside (DDM, ULTROL grade), dimethyl sulfoxide (DMSO, Molecular Biology grade), chlorpromazine hydrochloride (CPZ, ≥ 98% TLC) and fluvoxamine maleate (FLUV, ≥ 97% HPLC) were purchased from Merck.

All aqueous solutions were prepared using 18.2 MΩ ultra-pure water and filtered with sterile 0.22 μm polyethersulfone (PES) membranes.

### Peptides

Fusion peptide domains (FP1, FP2 and FP1-FP2) of the SARS-CoV-2 spike protein were synthesized by GenScript with a purity ≥ 95%. The produced sequence were FP1(816-SFIEDLLFNKVTLADAGF-833), FP2 (834-IKQYGDCLGDIAARDLICAQKF-855) and FP1-FP2 (816-SFIEDLLFNKVTLADAGFIKQYGDCLGDIAARDLICAQKF-855); the 3 constructs included a C-terminal Leu-His_6_ tag. Lyophilized samples (1 mg aliquots) were solubilized in 1 mL of buffer H (25 mM HEPES/KOH, pH 7.5; 150 mM KCl; 10% (v/v) glycerol) by vortexing for 2 min at room temperature. Samples were snap-frozen in liquid nitrogen and stored at -80°C as aliquots of 50 μL.

### Liposomes

Liposomes were generated by the detergent-assisted method ^29^. 0.9 μmol of the appropriate lipid mixtures in chloroform were dried in glass tubes for 10 min under a gentle stream of argon, followed by 2 hours under vacuum. The dried lipid films were resuspended in 300 μL of buffer H containing 1% (w/v) β-OG by vigorously vortexing for 30 min at room temperature or 50°C when CER was included in the lipid mixture. The detergent concentration was next reduced below the critical micellar concentration, 0.33% (w/v), by slowly adding 600 μL of buffer H, and then removed by overnight flow dialysis against 4 L of buffer H. Liposomes (final lipid concentration of 1 mM) were stored on ice and protected from light for up to 2-3 weeks.

### FRET-based lipid mixing assay

27 μL of acceptor (non-fluorescent) liposomes at 1 mM and 21 μL of buffer H (or 15 μL of buffer H and 6 μL of drugs at 500 μM in DMSO 10% (v/v) in buffer H for experiments performed in the presence of drugs) were added to a flat bottom 96-well white polystyrene plate (Thermo Scientific) and pre-warmed at 37°C for 7 min; 6 μL of donor (fluorescent) liposomes at 500 μM were carefully added to one side of the well; 6 μL of peptides at 250 μM in buffer H were added to another side of the well. The fusion reaction was initiated by shaking the plate in order to mix the three different solutions. Lipid mixing was measured by following fluorescence dequenching of the NBD probes from the donor liposomes resulting from their dilution into the acceptor liposomes. The NBD fluorescence was monitored at 1-min intervals for 90 min (excitation at 460 nm; emission at 535 nm; cutoff at 530 nm) by the SpectraMax M5 microplate reader (Molecular Devices) equilibrated to 37°C. After 90 min, 10 μL of 2.5% (w/v) DDM was added to completely dissolve the liposomes and thus measure the NBD fluorescence at infinite dilution, F_max_(NBD); the data were then normalized using the following equation that gives the percentage of NBD fluorescence increase at time t, %F(NBD, t):

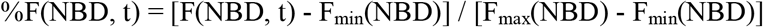

where F_min_(NBD) is the lowest NBD fluorescence value from all time points.

### Multi-angle dynamic light scattering

5 μL of liposomes at 1 mM and 95 μL of buffer H were mixed in a low volume quartz batch cuvette (ZEN2112, Malvern Panalytical) and their size distribution was determined at 37°C with the Zetasizer Ultra Red instrument (Malvern Panalytical) using the multi-angle dynamic light scattering (MADLS) mode, which measures the correlation function in three scattering directions: back scatter (173 degrees), side scatter (90 degrees) and forward scatter (13 degrees).

### Plasmids

The LgBit construct was obtained by removing the EGFP-Hsp70 sequence from pEGFP-Hsp70 plasmid (Addgene#15215), which was replaced by PCR-amplified LgBit insert from LgBit Expression Vector (Promega#N2681). AgeI and SpeI restriction enzymes (NEB) were used to digest both the plasmid and the insert before ligation using T4 DNA Ligase (NEB).

Similarly, the HiBit-Hsp70 construct was obtained by removing the EGFP-Hsp70 sequence from pEGFP-Hsp70 plasmid using AgeI and SpeI. The HiBit-Hsp70 insert was PCR-amplified directly from pEGFP-Hsp70 plasmid using primers containing the HiBit sequence and then inserted using AgeI and XbaI (NEB), which is SpeI compatible. The primers used to amplify LgBit sequence were: AgeI-LgBit forward 5’-TAG ACC GGT CAC CAT GGT CTT CAC ACT CGA AGA-3’ and SpeI-LgBit reverse 5’-TCC ACT AGT AAC GTT ACT CGG AAC AGC ATG GAG-3’. The primers used to amplify HiBit-Hsp70 sequence were: AgeI-HiBit-Hsp70 forward 5’-GCC ACC GGT ACC ATG ACT AGT GTG AGC GGC TGG CGG CTG TTC AAG AAG ATT AGC GGA TCC TCC GGT GGA TCG AGC GGT GGG AAT TCT GGT GGA GGA TCC GCT AGC ATG GCC AAA GCC GCG-3’ and XbaI-Hsp70 reverse 5’-GCA TCT AGA AGA GCT CGT CTC AAG CTT GCT AAT CTA CCT CCT CAA TGG TGGG-3’.

The plasmid of wild-type Spike with a C-terminal Flag tag was generated by the Biochemistry and Biophysics (B&B) facility of the Institute of Psychiatry and Neuroscience of Paris (IPNP). The vector pcDNA3.1(-) containing the SARS-CoV-2, Wuhan-Hu-1 Spike glycoprotein gene (pcDNA3.1-Spike-WT) was kindly provided by BEI resources (#NR-52420). The C-terminal Flag tag was introduced into pcDNA3.1-Spike-WT thanks to NEBuilder® HiFi DNA Assembly (NEB). The Flag tag was added during amplification of Spike-WT from pcDNA3.1-Spike-WT with the primers: forward 5’-ACC ACA AAG CGG ACA ATG TTC GTG TTT CTG GTG CTG-3’ and reverse 5’-ACC GAG CTC GGA TCC TCA TTT ATC ATC ATC ATC TTT ATA ATC GCT GCC GGT GTA GTG CAG CTT CACG-3’. The resulting plasmid called pcDNA3.1-Spike-WT-Flag was amplified and verified by sequencing.

The human ACE2 plasmid was purchased from VectorBuilder (#VB900122-0052prs).

### Cell culture

HeLa cells (ATCC) were cultured in DMEM (Gibco) complemented with 10% FBS (Gibco or Biosera) at 37°C and 5% CO_2_. Stable cell lines were selected with Geneticin 10 μg/mL (Gibco) after Lipofectamine 2000-based transfection (Invitrogen).

### Cell-cell fusion assay

100,000 donor cells (HiBit-Hsp70) were transfected or not with Spike-WT-Flag encoding plasmid (1 μg DNA / 1 μL Lipofectamine 2000) in a 24-well plate format. Similarly, 100,000 acceptor cells (LgBit) were transfected or not with ACE2 encoding plasmid (1 μg DNA / 1 μL Lipofectamine 2000). When required, donor and acceptor cells were only treated with Lipofectamine 2000 (mock). 8 hours post-transfection, cells were washed three times with PBS, trypsinized and resuspended in complete media. 20,000 cells (for each donor and/or acceptor cells) were seeded in a white 96-well plate format and cultured for 24 hours. When required, Fumonisin B1 (20 μM) or MβCD (2 mM), and/or CPZ (10 μM) were added for 24 hours prior to reading nanoluciferase activity, using Nano-Glo Live Cell Assay System (Promega) and SpectraMax iD3 microplate reader (Molecular Devices).

## Acknowledgments

D.T. is funded by the “Agence Nationale de la Recherche” (ANR-19-CE11-0018-01), the “Fondation de France – Recherche fondamentale et clinique sur les maladies psychiatriques” and the “Association Française contre les Myopathies” (AFM-Téléthon Research Grant 23778). Work in G.L laboratory is supported by INSERM, “Agence Nationale de la Recherche” (Excelldisc, BIOEV, EVfusion) and Chaire d’Excellence Idex, Université Paris Cité. Z.E.A.D. received a postdoctoral fellowship from “Fondation pour la Recherche Medicale” (FRM).

