## Supplementary Figures for "Cholesterol and ceramide facilitate SARS-CoV-2 Spike protein-mediated membrane fusion"

**Figure S1**

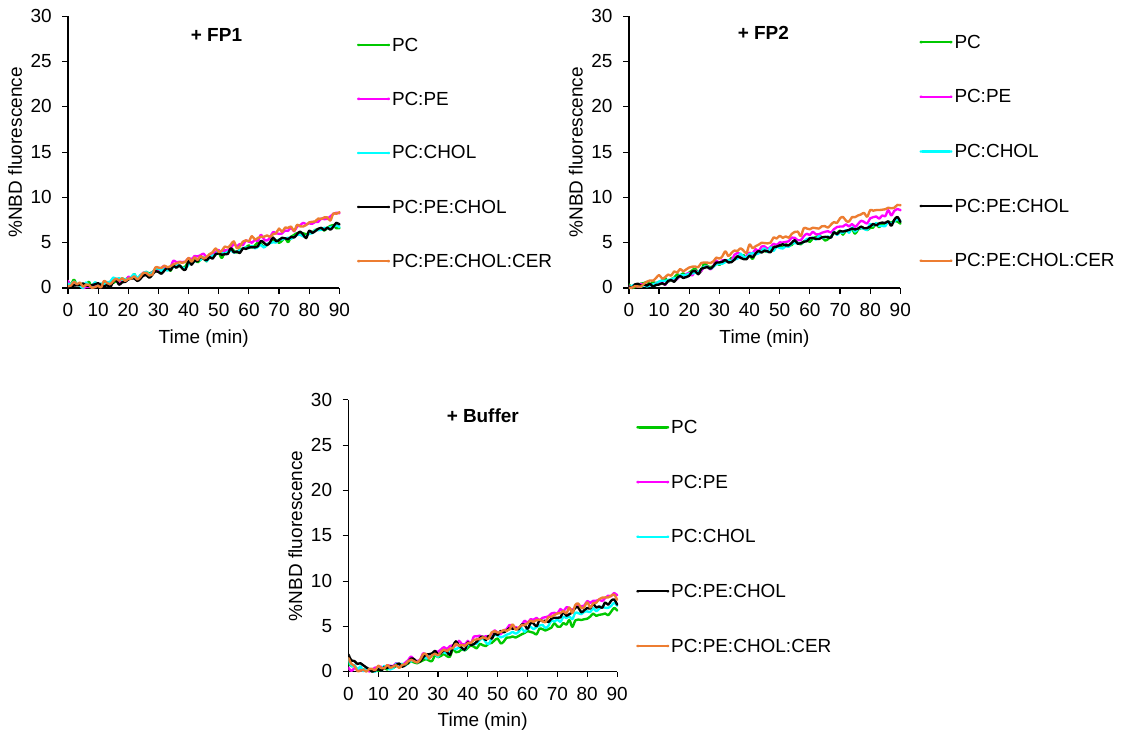

**Figure S1.** FRET-based lipid mixing experiments between PC:NBD:Rho (97:1.5:1.5) v-liposomes (lacking NTA-Ni lipid anchors) and PC (100), PC:PE (90:10), PC:CHOL (70:30), PC:PE:CHOL (60:10:30) or PC:PE:CHOL:CER (40:10:30:20) c-liposomes incubated at t=0 of the assay with FP1, FP2 or buffer H (500 µM of lipids and 25 µM of peptides). No significant lipid mixing was measured when FP1 was added to v-liposomes lacking NTA-Ni lipids, showing that only membrane-anchored FP1 has the capacity to induce fusion between v- and c-liposomes (Fig. 2).

**Figure S2**

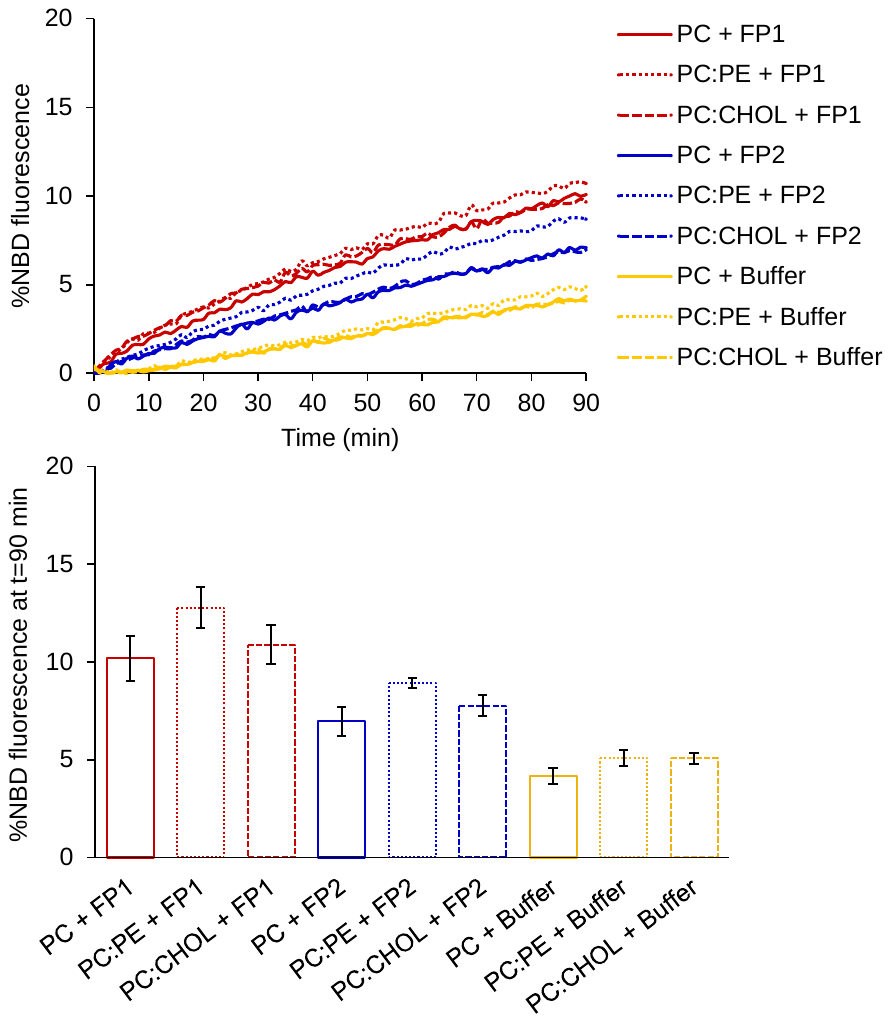

**Figure S2.** Fusion measured with the FRET-based lipid mixing assay between PC:NTA-Ni:NBD:Rho (92:5:1.5:1.5) v-liposomes and PC (100), PC:PE (90:10) or PC:CHOL (70:30) c-liposomes in the absence (yellow) or presence of FP1 (red) or FP2 (blue) added at t=0 (500 µM of lipids and 25 µM of peptides). Fusion was not significantly activated when the c-liposome membrane contained only PE or only CHOL (in addition to PC). The top panel shows one representative set of experiments, and the bottom panel the average extent of lipid mixing after 90 min of reaction (n=3-9 independent experiments; error bars are standard errors).

**Figure S3**

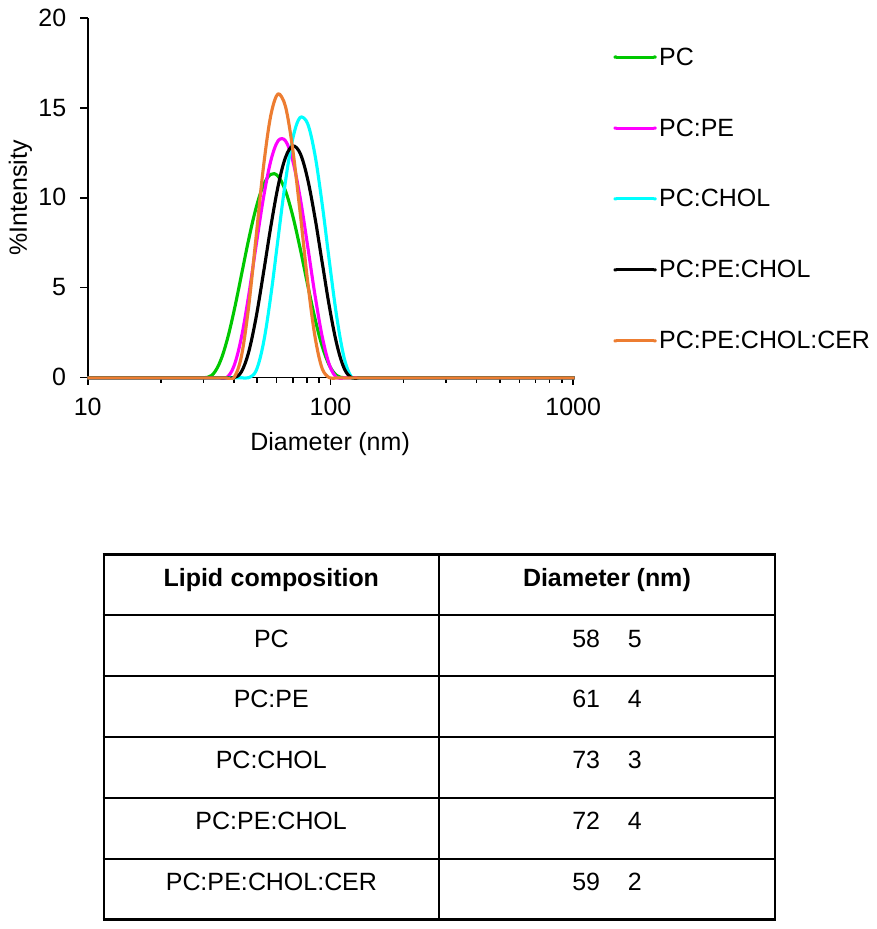

**Figure S3.** Liposome size distribution as a function of their lipid composition determined by multi-angle dynamic light scattering. PE, CHOL and CER lipids were added at 10, 30 and 20 mol% (at the expense of PC), respectively. The top panel shows one representative set of measurements, and the bottom panel the average position of the peak (n=3-6 independent liposome reconstitutions). The size of liposomes was not affected by their lipid composition. Importantly, higher FP1-mediated fusion observed with PC:PE:CHOL (60:10:30) and PC:PE:CHOL:CER (40:10:30:20) c-liposomes (Fig. 2) cannot be explained by their size since they are overall larger than pure PC c-liposomes (smaller, highly curved, liposomes are known to be more fusogenic[^1–3^](https://sciwheel.com/work/citation?ids=977817,13626619,13626620&pre=&pre=&pre=&suf=&suf=&suf=&sa=0,0,0&dbf=0&dbf=0&dbf=0)).

**Figure S4**

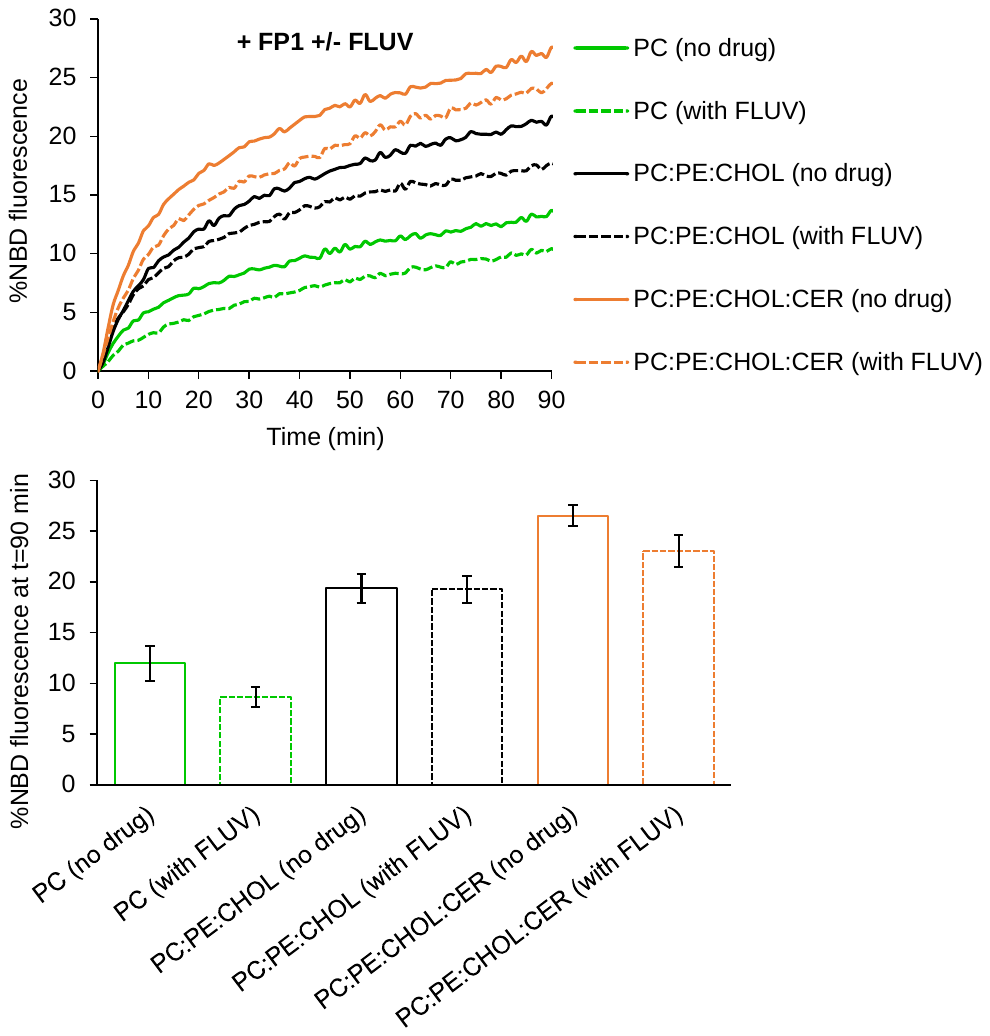

**Figure S4.** FRET-based lipid mixing experiments between PC:NTA-Ni:NBD:Rho (92:5:1.5:1.5) v-liposomes and PC (100), PC:PE:CHOL (60:10:30) or PC:PE:CHOL:CER (40:10:30:20) c-liposomes in the presence of FP1 and in the presence/absence of fluvoxamine (FLUV), both added at t=0 (500 µM of lipids, 25 µM of peptides and 50 µM of FLUV). All fusion experiments (with or without FLUV) were performed in a final DMSO concentration of 1% (v/v) in buffer H. FLUV slightly inhibited FP1-mediated fusion between v- and c-liposomes independently of the c-liposome membrane lipid composition. The top panel shows one representative set of experiments, and the bottom panel the average extent of lipid mixing after 90 min of reaction (n=3-6 independent experiments; error bars are standard errors).

**Figure S5**

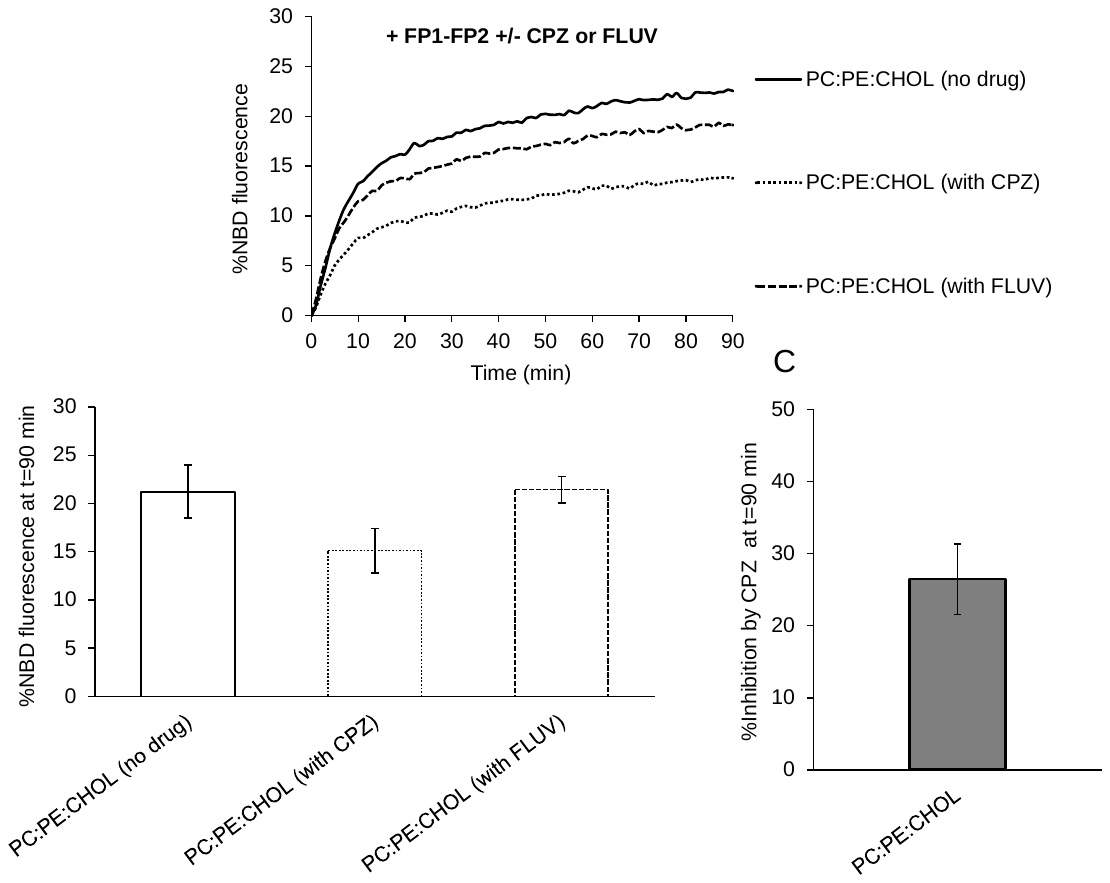

**Figure S5.** Fusion measured with the FRET-based lipid mixing assay between PC:NTA-Ni:NBD:Rho (92:5:1.5:1.5) v-liposomes and PC:PE:CHOL (60:10:30) c-liposomes incubated at t=0 of the assay with FP1-FP2 in the presence/absence of CPZ or FLUV (500 µM of lipids, 25 µM of peptides and 50 µM of CPZ or FLUV). FP1-FP2 induced robust lipid mixing between v- and c-liposomes, and CPZ (but not FLUV) strongly inhibited FP1-FP2-mediated liposome fusion. The extent of lipid mixing with FP1-FP2 and the level of fusion inhibition by CPZ after 90 min are similar to those measured with FP1 (Fig. 3). The top panel shows one representative set of experiments. The bottom panels show the average extent of lipid mixing and the percentage of fusion inhibition by CPZ after 90 min (n=3-4 independent experiments; error bars are standard errors).
